# Characterization of proteoform post-translational modifications by top-down and bottom-up mass spectrometry in conjunction with UniProt annotations

**DOI:** 10.1101/2023.04.04.535618

**Authors:** Wenrong Chen, Zhengming Ding, Yong Zang, Xiaowen Liu

## Abstract

Many proteoforms can be produced from a gene due to genetic mutations, alternative splicing, post-translational modifications (PTMs), and other variations. PTMs in proteoforms play critical roles in cell signaling, protein degradation, and other biological processes. Mass spectrometry (MS) is the primary technique for investigating PTMs in proteoforms, and two alternative MS approaches, top-down and bottom-up, have complementary strengths. The combination of the two approaches has the potential to increase the sensitivity and accuracy in PTM identification and characterization. In addition, protein and PTM knowledgebases, such as UniProt, provide valuable information for PTM characterization and validation. Here, we present a software pipeline called PTM-TBA (PTM characterization by Top-down, Bottom-up MS and Annotations) for identifying and localizing PTMs in proteoforms by integrating top-down and bottom-up MS as well as UniProt annotations. We identified 1,662 mass shifts from a top-down MS data set of SW480 cells, 545 (33%) of which were matched to 12 common PTMs, and 351 of which were localized. PTM-TBA validated 346 of the 1,662 mass shifts using UniProt annotations or a bottom-up MS data set of SW480 cells.

## 1. Introduction

Post-translational modifications (PTMs) in proteoforms, like methylation, acetylation, and phosphorylation, play crucial biological roles in biological systems and diseases [1, 2]. For example, kinase phosphorylation is essential for signal transduction in cells and the development of cancer cells [3]. PTM characterization, which identifies and localizes PTMs in proteoforms, is important for studying protein functions, understanding biological mechanisms, discovering disease biomarkers, and designing personalized vaccines [2, 4-7].

The dominant techniques for identifying PTMs in proteoforms are two complementary mass spectrometry (MS) approaches: bottom-up and top-down MS [5, 8, 9]. In bottom-up MS [5], proteoforms are proteolytically digested into short peptides, which are separated by liquid chromatography (LC) or other methods and then analyzed by MS. Bottom-up MS usually provides high fragment ion coverage of identified peptides, which increases the accuracy in determining PTM types and locations in peptides. However, bottom-up MS may identify only several peptides of a protein and miss many peptides with PTMs in proteome-wide analyses [10]. And PTM combinatorial patterns in proteoforms are lost in digestion, making bottom-up MS inefficient for analyzing complex proteoforms with multiple PTMs [11]. Top-down MS analyzes intact proteins instead of peptides [12], so it can identify PTM combinatorial patterns in proteoforms and characterize proteoforms with multiple PTMs. But top-down MS still suffers from limited sensitivity and throughput and often fails to identify low-abundance proteoforms in proteome-wide studies. Top-down mass spectra also tend to miss many fragment ions, resulting in ambiguous localization sites for PTMs [13].

Continuous efforts have been made to develop computational methods for identifying and localizing PTMs using bottom-up MS [14-18]. These methods identify PTMs in peptides by database search with pre-specified variable PTMs [14, 16, 17] or open search [10, 15, 18, 19]. While database search with variable PTMs reports the types of identified PTMs directly, the open search method first identifies unexpected mass shifts, which are then matched common PTMs to identify their PTM types. These PTMs are further localized to determine their modification sites [20]. Many software tools, such as MaxQuant [14], determine the site of a PTM in an identified peptide-spectrum-match (PSM) based on the similarity scores between the spectrum and all candidate forms of the peptide with the PTM on different sites. The modified peptide form with the best similarity score is reported as the PTM localization result. The confidence scores of PTM characterization or localization results are computed using probabilistic or machine learning models, such as AScore [20], SLoMo [21], PhosphoRS [22] and Andromeda score [23]. Similarly, in top-down MS, PTMs are identified by database search with variable PTMs [24, 25], the open search strategy [26, 27], or spectral alignment [27]. Confidence scores of PTM characterization and localization results are reported using Bayesian or other statistical models [26, 28, 29].

Combining bottom-up and top-down MS is a promising direction for PTM identification and characterization because the two approaches have complementary strengths [11, 30]. Bottom-up mass spectra provide high fragment ion coverage that top-down mass spectra often lack, while top-down mass spectra offer PTM combinatorial pattern information for characterizing complex proteoforms. One approach can also validate PTMs identified by the other. Several existing tools, like Proteoform Suite [30], adopt this method for PTM characterization.

Protein and PTM knowledgebases offer additional evidence for validating PTMs identified by MS [31]. Protein annotations in UniProt [32] contain many PTMs reported in the literature. Additionally, dbPTM [33], SysPTM [34], and PRISMOID [35] store both structural and functional information about PTMs.

Here, we present PTM-TBA, a software pipeline for proteoform PTM characterization by combining top-down MS, bottom-up MS, and UniProt PTM annotations. We performed a systematic evaluation of the pipeline using top-down MS, bottom-up MS data of SW480 colorectal cancer cells. Using top-down MS, we identified 1,662 mass shifts in proteoforms reported from SW480 cells, of which 545 were matched to 12 high frequency PTMs, and 351 were confidently localized. We also validated 334 PTM sites using peptides identified by bottom-up MS data and/or UniProt annotations.

## 2. Methods

### 2.1 Data sets

A bottom-up MS data set and a top-down MS data set of SW480 colorectal cancer cells were used to evaluate the proposed PTM characterization and validating pipeline. The bottom-up MS experiment [36] was performed in technical triplicate, and only the first replicate was used in this paper. Proteins of SW480 cells were digested using trypsin. The dried peptides were fractionated into five fractions using 8%, 15%, 22%, 30%, and 50% ACN in 10nM TEAB (pH 9). A Waters NanoAcquity LC system with a BEH C18 column (Waters, 10 cm × 100 mm, 1.7 μm particle size) coupled with a Q-Exactive mass spectrometer (Thermo Fisher) was used in the analysis. The sample peptides were separated over a 90-min linear gradient (A: 0.1% formic acid in water, B: 0.1% formic acid in acetonitrile). The gradient was used for the samples with solvent B added from 2% to 80% from 0 to 80 min and re-equilibrated at 2% from 80 to 90 min. MS1 scans were collected from 350 to 2,000 m/z at a resolution of 70,000 (at 200 m/z) with an AGC target of 1 × 10^6^ ions, and MS/MS scans were collected from 100 to 1,500 m/z at a resolution of 17,500 (at 200 m/z) with an AGC target of 5 × 10^5^ ions. The top 12 precursor ions in each MS1 spectrum were isolated with a 2 m/z window for MS/MS analyses and the normalized collision energy was set to 28.

The top-down MS data set of SW480 cells was generated using size exclusion chromatography (SEC)-capillary zone electrophoresis (CZE)-MS/MS [37]. The sample proteins were initially separated into 6 fractions using an SEC column, and each fraction was then injected into a fused silica capillary with a linear polyacrylamide (LPA) coating and a background electrolyte of 5% acetic acid for a 100-min separation. The electrospray voltage was set to between 2.2 and 2.3 kV, and the separation voltage was 30 kV. The CZE system was connected to a Q-Exactive HF mass spectrometer (Thermo Fisher) for MS/MS analysis. The resolution of the MS1 and MS/MS spectra was 120,000 at 200 m/z. Using higher-energy C-trap dissociation (HCD) MS/MS, the top 5 precursor ions in each MS1 spectra were fragmented. Three technical replicates with a total of 18 runs (6 fractions × 3 replicates) were obtained for SW480 cells, and only the first replicate was used in this paper.

### 2.2 Bottom-up MS database search

All raw files were converted to centroided mzML files using msconvert in ProteoWizard [38]. A human proteome database, GENCODE-UniProt (18,417 proteins), was built based on the protein sequences shared by the GENCODE basic annotation (version 38, 19,652 proteins) [39] and the UniProt human proteome database (version 11/23/2019, 20,350 proteins) [32] using TopPG (version 1.0) [40]. It contained a reference protein sequence for each gene that was annotated by both UniProt and the GENCODE basic annotation. The centroided bottom-up mass spectra were searched against the GENCODE-UniProt database concatenated with a reversed decoy database (18,417 entries) using MSFragger [15] (version 3.4) with the open search strategy. Methionine oxidation and N-terminal acetylation were chosen as variable PTMs, cysteine carbamidomethylation as the fixed modification. Up to 3 variable modifications were allowed in each identified peptide, and the default setting [-150 Da, 500 Da] was used for the allowed mass shift of precursor masses. Identified PSMs were filtered using a 1% spectrum-level false discovery rate (FDR). Then the reported PSMs were grouped to obtain peptide identifications, which were filtered using a 1% peptide-level FDR. Finally, identified proteins were filtered with a 1% protein-level FDR.

### 2.3 Top-down MS data analysis

TopFD (version 1.5.2) [41] was employed to deconvolute the centroided top-down mass spectra to neutral monoisotopic masses of precursor and fragment ions. The deconvoluted MS/MS spectra were searched against the GENCODE-SWISS database concatenated with a decoy database with the same size using TopPIC (version 1.5.2) [27]. The parameter settings used in database search were set to the following: an error tolerance of 15 ppm for precursor and fragment masses, at most one unexpected mass shift in each identified proteoform, and cysteine carbamidomethylation as the fixed modification. Identified proteoforms were filtered with a 1% proteoform-level FDR.

### 2.4 Matching mass shifts identified in proteoforms and peptides

Mass shifts identified in a peptide/proteoform were matched to those in another peptide/proteoform for removing duplicated shifts and validating identifications. A mass shift and its possible sites in a peptide/proteoform are represented by a quadruple [*m, p, a, b*], where *m* is the mass shift, *p* is the protein containing it, and positions *a* and *b* specify a region [*a, b*] of the protein that contains potential modification sites of the mass shift. In top-down MS, the region [*a, b*] of a mass shift reported by TopPIC was slightly extended to correct possible errors in [*a, b*] introduced by randomly matched fragment peaks. Specifically, position *a* was extended to the left until the extended part contained 2 matched fragment ions or the N-terminus was reached, and position *b* was extended to the right similarly. When the PTM type of a mass shift is known, the mass shift with its PTM type is represented by a quintuple [*m, p, a*, b, *t*], where *t* is the type of the PTM.

A mass shift [*m*_1_, *p*_1_, *a*_1_, *b*_1_] is matched to another mass shift [*m*_2_, *p*_2_, *a*_2_, *b*_2_] if (1) *p*_1_ and *p*_2_ are the same, (2) the sequence regions [*a*_1_, *b*_1_] and [*a*_2_, *b*_2_] overlap, and (3) the minimum difference among the mass pairs (*m*_1_, *m*_2_), (*m*_1_, *m*_2_-1.00235), (*m*_1_, *m*_2_+1.00235) is smaller than an error tolerance (0.1 Da in the experiments). The mass difference of 1.00235 Da is allowed in the comparison because it is a common error in deconvoluted precursor masses in top-down MS. A mass shift with its PTM type [*m*_1_, *p*_1_, *a*_1_, *b*_1_, *t*_1_] is matched to another mass shift with its PTM type [*m*_2_, *p*_2_, *a*_2_, *b*_2_, *t*_2_] in a proteoform if (1) the mass shift [*m*_1_, *p*_1_, *a*_1_, *b*_1_] is matched to [*m*_2_, *p*_2_, *a*_2_, *b*_2_], (2) *t*_1_ and *t*_2_ are the same, and (3) the overlapping region of [*a*_1_, *b*_1_] and [*a*_2_, *b*_2_] contains at least one amino acid residue that can be modified by the PTM.

### 2.6 PTMs in UniProt protein annotations

An annotation file of the UniProt human proteome (version 06/16/2022, 204, 906 entries) was downloaded from UniProt [32]. PTMs and their sites in proteins were extracted from the annotation file using a Python script in PTM-TBA. A PTM annotation is matched to a mass shift [*m, p, a, b*] in a proteoform if the annotated PTM is in the region [*a, b*] of protein *p* and the mass shift of the PTM is matched to the mass shift *m* with an error tolerance (0.1 Da in the experiment). The error +/-1.00235 Da is also allowed in the matching of the mass shift.

### 2.7 Removing duplicated mass shifts

To remove duplicated mass shifts, we first grouped mass shifts reported from top-down or bottom-up MS data into clusters and then removed duplicated mass shifts in each cluster. In the clustering step, two mass shifts [*m*_1_, *p*_1_, *a*_1_, *b*_1_] and [*m*_2_, *p*_2_, *a*_2_, *b*_2_] are added to the same cluster if *p*_1_ and *p*_2_ are the same and the difference between *m*_1_ and *m*_2_ is smaller than an error tolerance (0.1 Da in the experiments). To remove duplicated mass shifts in a cluster, the mass shifts in the cluster are ranked using the left boundary (*a* in the quadruple representation [*m, p, a, b*]) of the range containing possible modification sites. Next, we iteratively check the mass shifts in the cluster following their ranks to remove duplicated ones using a greedy algorithm (Fig. S1 in the supplemental material).

## 3. Results

### 3.1 Overview of the PTM identification and localization pipeline

Figure 1 illustrates the overall scheme of the PTM-TBA pipeline for PTM identification, localization, and validation using top-down MS, bottom-up MS data, and UniProt annotations. Top-down MS and bottom-up MS data were generated from SW480 cells separately. MSFragger [15] was employed to identify mass shifts in peptides by database search using bottom-up MS data. After duplicated shifts were removed, mass shifts in peptides were divided into the four classes: Class I: N-terminal acetylation; Class II: high frequency PTMs with localized PTM sites (excluding N-terminal acetylation); Class III: high frequency PTMs without PTM localization; and Class IV: low frequency mass shifts. TopPIC [27] was used to identified mass shifts from top-down MS. After duplicated shifts were removed (Methods), the identified mass shifts were also divided into four classes. The mass shifts in proteoforms were validated by matching them to mass shifts in peptides. Human protein PTM annotations were extracted from the UniProt knowledgebase. Mass shifts in proteoforms were further validated by matching these PTM annotations to the mass shifts in proteoforms.

**Figure 1.**
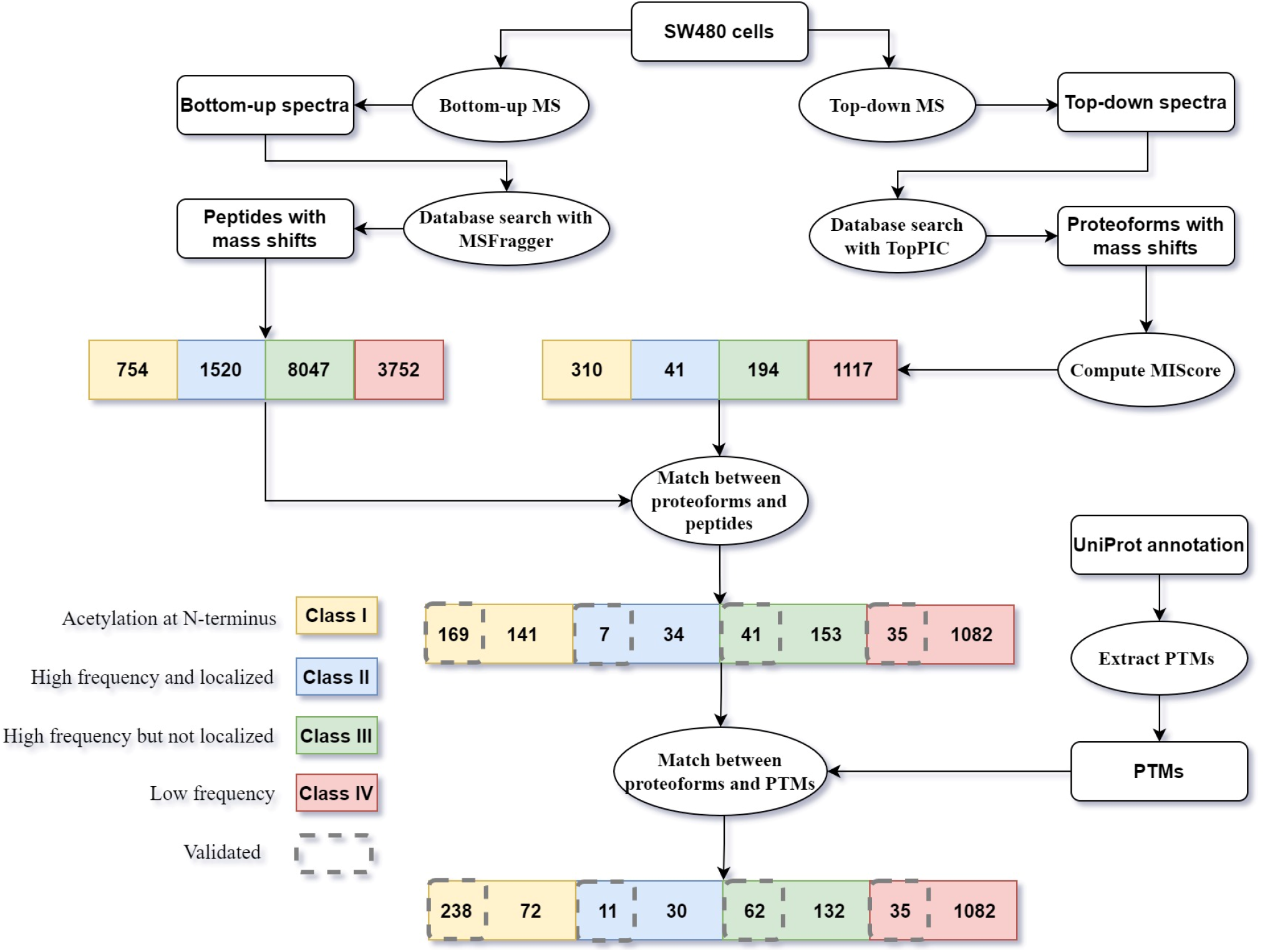
The overall scheme of the pipeline for proteoform PTM identification, localization, and validation using top-down MS, bottom-up MS and UniProt annotations.

### 3.2 Mass shift identification by bottom-up MS

TopPG [40] was employed to generate a human protein database GENCODE-UniProt (18,417 proteins), which contained protein sequences shared by the GENCODE basic annotation and the UniProt human proteome database (Methods). A total of 99,427 MS/MS spectra were obtained from the bottom-up MS experiment of SW480 cells, which were searched against the GENCODE-UniProt database for peptide identification using MSFragger [15] (Methods). Identified peptide-spectrum-matches (PRMs), peptides, proteins were filtered at a 1% PSM-level, peptide-level, and protein-level FDR, respectively. As a result, 72,241 PSMs, 28,141 peptides, and 3,825 proteins were reported after filtering at 1% PSM-level, peptide-level, and protein-level FDR. Of the 72,241 identified PSMs, 52,845 were from unmodified peptides and the remaining 19,396 were from modified peptides, which contained a total of 24,341 mass shifts (some identified peptides contained more than one mass shift/PTM.

A mass shift in a peptide may be reported in several identified PSMs (Fig. 2(a)), so duplicated mass shifts need to be removed (Methods). After duplicated mass shifts were removed, 14,073 mass shifts/PTMs sites were reported, including 754 N-terminal acetylation sites and 9567 mass shifts that were matched to the shift of a PTM. A total of 12 PTMs (Table S1 in the supplemental material) were identified with a high frequency (observed in > 0.15% of all identified PSMs). Fig.3 shows the frequencies of mass shifts in the range [0, 200] Da, in which high frequency PTMs were labeled. Using the 12 high frequency PTMs, the 14,073 mass shifts were divided into the four classes: 754 class I, 1,520 class II, 8,047 class III, and 3,752 class IV mass shifts.

**Figure 2.**
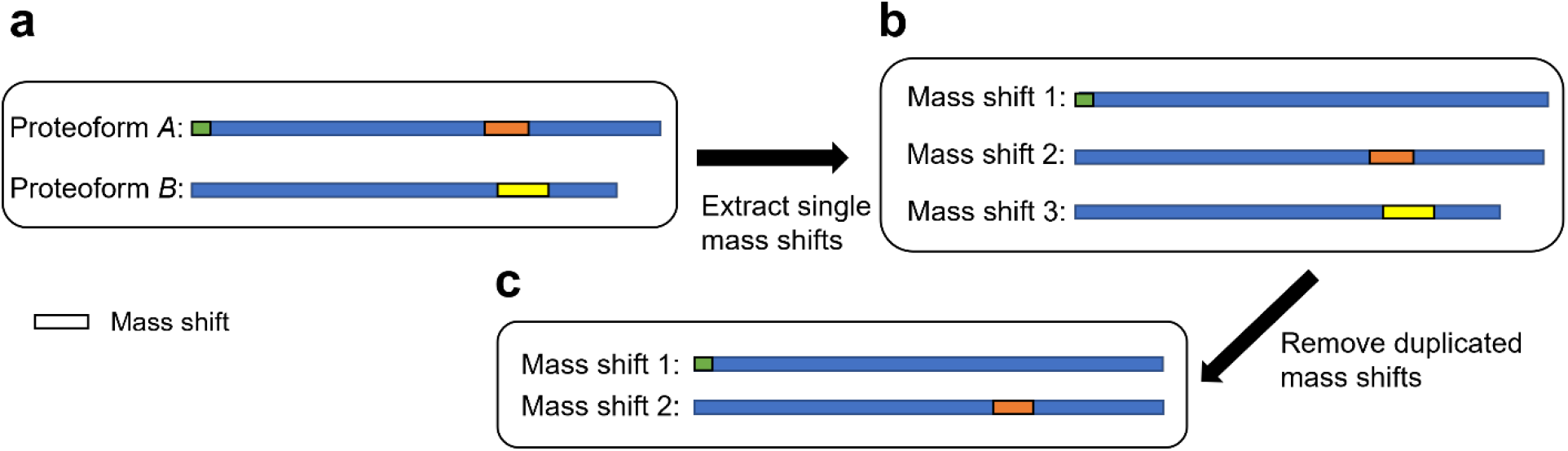
An illustration of the extraction and duplication removal of mass shifts in proteoforms. (a) Proteoform identifications A and B are from the same protein. Proteoform A contains the whole protein sequence with two mass shifts (green and brown) and proteoform B is a truncated one with one mass shift (yellow). The colored parts show protein regions containing possible modification sites of the mass shift. (b) Three single mass shifts are extracted from the two proteoform identifications. (c) The brown and yellow mass shifts have similar shifts and their possible modification site regions overlap, so they are treated as duplicated ones and only one is kept.

**Figure 3.**
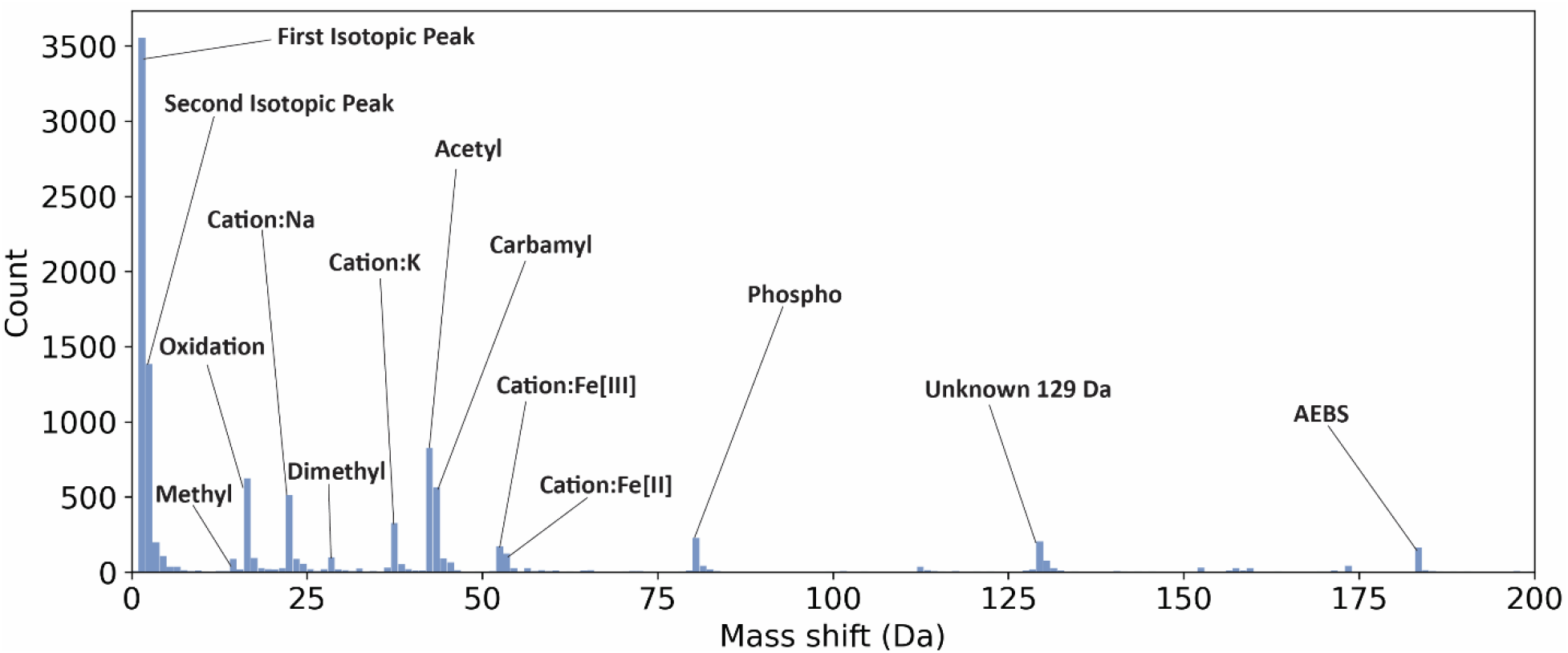
A histogram of mass shifts reported by MSFragger from the bottom-up MS data in the range [0, 200] Da.

### 3.3 Mass shifts identified in top-down MS

The top-down MS data of SW480 cells contained 22,455 MS/MS spectra, which were searched against the GENCODE-UniProt database using TopPIC [27] (version 1.5.2) (Methods). TopPIC reported 2,153 proteoforms with unexpected mass shifts and/or N-terminal acetylation, including 69 histone proteoforms, and 1,494 proteoforms without any mass shifts/PTMs except for cysteine carbamidomethylation from 828 proteins with a 1% proteoform-level FDR. The 69 histone proteoforms were excluded from downstream analysis because most of them contained multiple PTMs.

Similar to mass shifts in peptides, duplicated mass shifts and PTMs in identified proteoforms were removed. After removing duplicated mass shifts, we identified 310 N-terminal acetylation sites (Supplemental Table S2) and 1,352 other mass shifts in the 1,613 proteoform, of which 59 contained both N-terminal acetylation and another mass shift. Fig. 4 shows the distribution of the identified mass shifts in [0, 200] Da. The distribution of mass shifts in [-500, 500] Da is given in Fig. S2 in the Supplemental Material. Oxidation, methylation, acetylation, and phosphorylation are the most frequently observed PTMs, which are consistent with those reported by bottom-up MS. Additionally, 134 mass shifts (9.91% of all reported mass shifts) have a shift around -43 Da, which can be explained by an incorrect assignment of carbamidomethylation on a cysteine with methylation.

**Figure 4.**
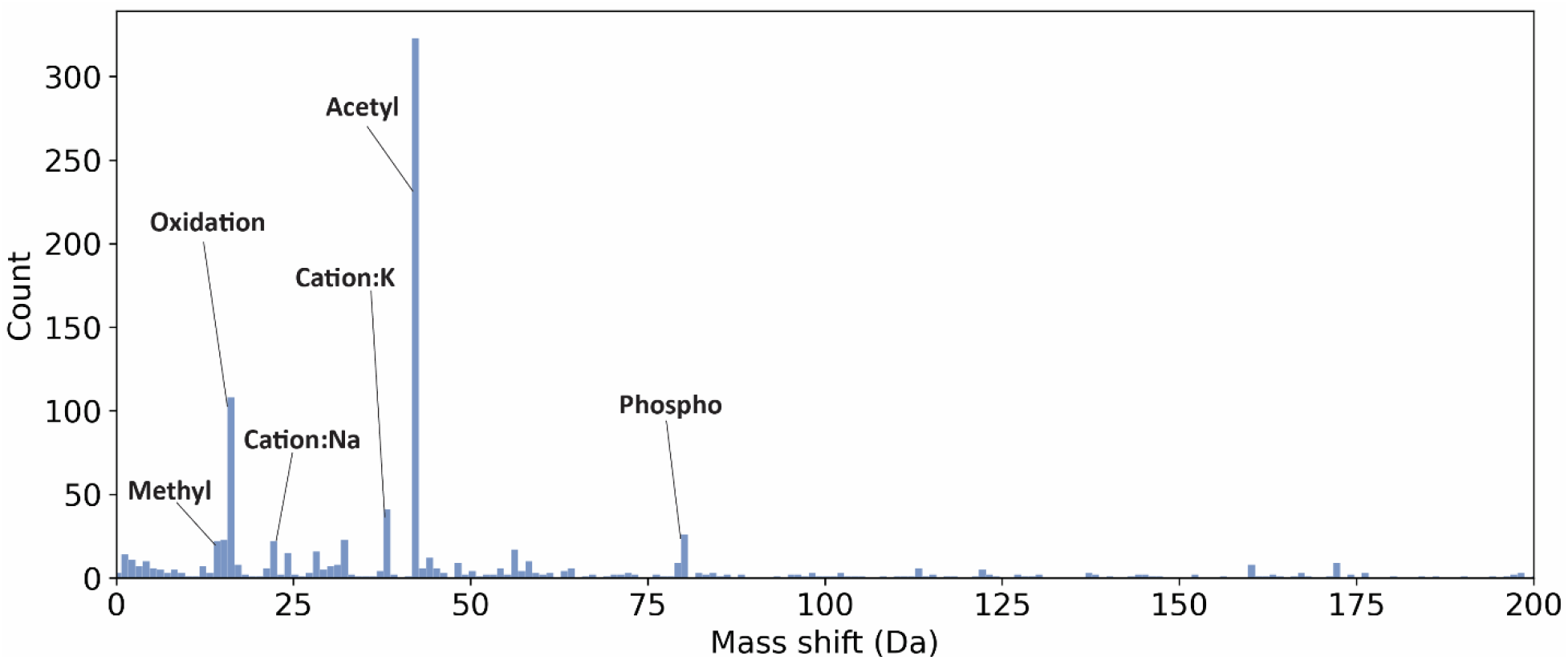
A histogram of mass shifts reported by TopPIC from the top-down MS data in the range [0, 200] Da.

The 1,352 mass shifts were further matched to high frequency PTMs identified from bottom-up MS to determine their PTM types. Because the mass shift (0.98 Da) of deamidation is similar to +1 Dalton errors in deconvoluted precursor masses, which are commonly observed, deamidation was excluded and only the remaining 11 high frequency PTMs in Supplemental Table 1 were matched to the mass shifts reported by top-down MS. The PTM type of a mass shift was determined if (1) the mass shift matched the shift of a high frequency PTM with an error tolerance of 0.1 Da (The error +/-1.00235 Da was also allowed in the matching of the mass shift) and (2) the possible site of the mass shift contained at least one amino acid that can be modified by the PTM. Using the method, a total of 235 mass shifts were identified as high frequency PTMs (Supplemental Table S2), including 194 oxidations. For these 235 high frequency PTMs, MIScore [29] was used to localize their PTM sites, and 41 PTMs were confidently localized with a confidence score ≥0.6 (Supplemental Table S2).

As a result, all the identified N-terminal acetylation sites and unexpected mass shifts were divided into 4 classes: (I) 310 N-terminal acetylation sites, (II) 41 high frequency PTMs with localized sites (excluding N-terminal acetylation), (III) 194 high frequency PTMs without localization, and (IV) 1,117 low frequency mass shifts.

### 3.4 Mass shift and PTM validation by bottom-up MS identification

The 1,117 Class IV mass shifts identified by top-down MS were compared with all mass shifts (14,073 entries) reported from bottom-up MS by MSFragger, resulting in 35 matched mass shifts between identified proteoforms and peptides. Of the 35 matched mass shifts, 27 were matched to a PTM in the UNIMOD database (version 10-17-2019) [42] and the others were not matched any PTMs in UNIMOD. The N-terminal acetylation sites in Class I and other high frequency PTMs in Class II and III reported by top-down MS were then compared with the high frequency PTMs in peptides reported by MSFragger (Methods), and 169 Class I, 7 Class II, 41 Class III PTMs were matched to those in peptides (Fig. 1).

### 3.5 PTM validation using UniProt annotations

The Class I, II, III mass shifts in proteoforms were compared with all PTM annotations extracted from the UniProt database (Methods) and 220 Class I, 8 Class II, and 33 Class III mass shifts were validated by matching them to UniProt annotations. The 42 validated mass shifts in Classes II and III included 8 methylation, 1 oxidation, 6 di-methylation, 3 acetylation, and 24 phosphorylation sites. Fig. 5(a) shows the numbers of mass shifts for the 4 classes validated by bottom-up MS only, UniProt annotations only, or both. Bottom-up MS and UniProt annotations provide complementary information for PTM validation. Of the 346 PTMs validated by bottom-up MS or UniProt annotations, 94 PTMs (27.1%) were validated by UniProt annotations only and 85 (24.6%) by bottom-up MS only.

**Figure 5.**
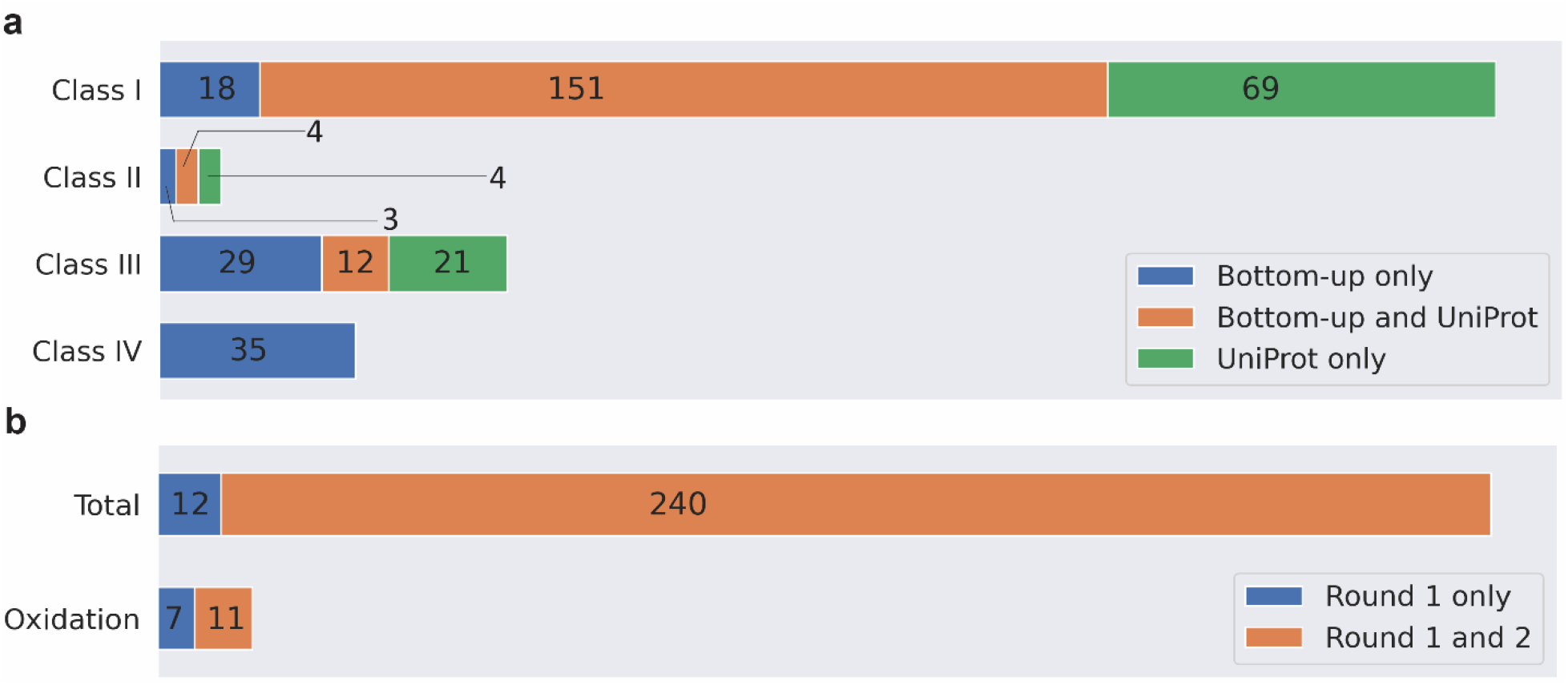
(a) Comparison of mass shifts in to-down proteoform identifications validated by bottom-up MS and UniProt annotations in the four classes. (b) Comparison of mass shifts (total and oxidation) validated in two rounds of MSFragger database search. In round 1, N-terminus acetylation and methionine oxidation were set as variable PTMs. In round 2, only N-terminus acetylation was set as the variable PTM.

### 3.6 Variable PTMs in bottom-up MS database search

The sensitivity of PTM identification can be improved by setting common PTMs as variable PTMs in database search. Methionine oxidation was chosen as a variable PTM in the previous analysis to increase the number of oxidation identifications in database search of the bottom-up MS data. To evaluate PTM identification without variable PTMs, we reanalyzed the bottom-up MS data using MSFragger by setting only N-terminal acetylation as the variable PTM. All other parameters were the same as the previous analysis.

The second round of bottom-up MS database search identified 72,199 PSMs, 28,169 peptides, and 3,841 proteins with a 1% FDR, which were similar to the numbers of identifications in the first round. We extracted 23,080 mass shifts from 19,264 identified PSMs with mass shifts or PTMs and finally obtained 13,602 mass shifts after removing duplicated ones. These mass shifts can be divided into the four classes: Class I: 764, Class II: 1,588, Class III 8,117, and Class IV: 3,133 (Fig. S2). When methionine oxidation was not set as a variable PTM, MSFragger identified only 274 oxidation sites, about 40.9% of the methionine oxidation sites (670) reported in the previous analysis, showing that setting a common PTM as a variable PTM can significantly increase the number of PTM sites identified by MSFragger. Besides of methionine oxidation, MSFragger reported similar numbers of modified sites for other mass shifts with the two different settings of variable PTMs. (Fig. 5(b)).

Similarly, the 13,602 mass shifts reported by the second-round database search of MSFragger validated 36 Class IV mass shifts in proteoforms reported by top-down MS data, of which 24 were matched to UNIMOD PTMs and 8 were not. Additionally, the 167, 7, 33 Class I, II, III mass shifts in proteoforms can be validated by mass shifts in peptides, respectively. Of the 252 mass shifts (including N-terminal acetylation) in proteoforms identified by top-down MS and validated by bottom-up MS in round one, 240 (95.2%) were also validated in round 2 (Fig. 5). The number of validated oxidation sites were reduced from 18 to 11 in the second round, showing that choosing appropriate variable PTMs is important for identifying PTMs in bottom-up MS (Fig. 5).

## 4. Conclusions

We developed and evaluated PTM-TBA, a software pipeline for proteoform PTM identification, localization, and validation using top-down MS, bottom-up MS, and UniProt annotations. The pipeline successfully identified 545 (32.4% of all reported mass shifts) high frequency PTMs and localized 351 sites in proteoforms from SW480 cells by top-down MS. Besides, we validated 311 PTMs in proteoforms using peptides identified by bottom-up mass spectra and UniProt annotations. In addition, 35 uncommon mass shifts were identified by both top-down and bottom-up MS.

PTM-TBA facilitates the discovery of novel PTMs and identification of known PTMs and their combinations. Nevertheless, there are still many challenging problems in proteoform PTM identification and localization. First, proteoforms may have multiple PTMs with complex combinatorial patterns, which complicate PTM identification and localization. Second, it is difficult to identify PTMs in low abundance proteoforms. PTM-specified enrichment techniques can contribute to detecting PTMs in low abundance proteoforms/peptides. For example, TiO_2_ columns [43] or antibodies [44] can be used to enrich phosphopeptides for MS/MS analysis.

**Table 1.** PTMs identified by matching mass shifts reported in top-down proteoforms identifications to 11 PTMs with high frequency observed in bottom-up MS data.

| UNIMOD ID | PTM name | Monoisotopic mass (Da) | Modified residues | # Proteoforms |
| --- | --- | --- | --- | --- |
| 1 | Acetylation | 42.010565 | CKST | 7 |
| 5 | Carbamylation | 43.005814 | CKMRSTY | 4 |
| 21 | Phosphorylation | 79.966331 | STY | 29 |
| 30 | Cation: Na | 21.981943 | DE | 10 |
| 34 | Methylation | 14.015650 | CHKNQRILDEST | 19 |
| 35 | Oxidation | 15.994915 | CPKDNRY | 116 |
| 36 | Di-methylation | 28.0313 | KNR | 17 |
| 276 | Aminoethylbenzenesulfonylation | 183.035399 | HKSY | 1 |
| 530 | Cation: K | 37.95582 | DE | 26 |
| 952 | Cation: Fe [II] | 53.919289 | DE | 4 |
| 1870 | Cation: Fe [III] | 52.911464 | DE | 2 |

## Acknowledgments

The research was funded by NIH through the grants R01GM118470, R01CA247863, and R01AI141625.

## Code availability

The code of PTM-TBA is available at https://github.com/wenronchen/PTM-TBA.

## SUPPORTING INFORMATION

Supplemental Material 1: An algorithm for removing duplicated mass shifts

Figure S1: A greedy algorithm for removing duplicated mass shifts

Figure S2: A histogram of mass shifts reported by TopPIC from the top-down MS data in the range [-500, 500] Da

Table S1: High frequency PTMs reported by MSFragger in database search Table S2: Mass shifts reported in top-down MS.

Table S3: Mass shifts from top-down MS which were matched with UniProt annotations

Table S4: Mass shifts from top-down MS which were matched with mass shifts from bottom-up MS.

