## Supplemental Material for "Characterization of proteoform post-translational modifications by top-down and bottom-up mass spectrometry in conjunction with UniProt annotations"

[1]Department of BioHealth Informatics, Indiana University-Purdue University Indianapolis, Indianapolis, IN 46202, USA, [2]Department of Computer Science, Tulane School of Science and Engineering, Tulane University, New Orleans, LA 70118, USA, [3]Department of Biostatics and Health Data Sciences, Indiana University School of Medicine, Indianapolis, IN 46202, USA, [4]Center for Computational Biology and Bioinformatics, Indiana University School of Medicine, Indianapolis, IN 46202, USA [5]Tulane Center for Biomedical Informatics and Genomics, Tulane University, New Orleans, LA 70112, USA, [6]Deming Department of Medicine, Tulane University, New Orleans, LA 70112, USA

### 1. An algorithm for removing duplicated mass shifts

To remove duplicated mass shifts, we first group mass shifts reported from top-down or bottom-up MS data into clusters and then remove duplicated mass shifts in each cluster. In the clustering step, two mass shifts [$m_1$, $p_1$, $a_1$, $b_1$] and [$m_2$, $p_2$, $a_2$, $b_2$] are added to the same cluster if $p_1$ and $p_2$ are the same and the difference between $m_1$ and $m_2$ is smaller than an error tolerance. A greedy algorithm is used to remove duplicated mass shifts in the same cluster with the objective of reporting a set of non-duplicated mass shifts and maximizing the number of mass shifts (Fig. S1). We sort all mass shifts in a cluster from a protein in the increasing order of the left boundary. Let $L = S_1, S_2, \ldots, S_n$ be the sorted mass shifts of the cluster. We compare the boundaries ($a_1$, $b_1$) of mass shift $S_1$ with the boundaries ($a_2$, $b_2$) of $S_2$ to remove duplicated ones. There are three cases. Case 1: $b_1 \leq a_2$, that is, $S_1$ and $S_2$ do not overlap (Step 4 in Fig. S1). In this case, $S_1$ is removed from the mass shift list $L$ and added to the result list $R$. Case 2: $a_2 < b_1 < b_2$, that is, $S_1$ and $S_2$ partially overlap (Step 6 in Fig. S1). In this case, $S_2$ is removed from the list $L$. Case 3, $b_1 \geq b_2$, that is, $S_1$ fully covers $S_2$ (Step 8 in Fig. S1). In this case, $S_1$ is removed from the list $L$. The comparison step is repeated for the first two mass shifts in $L$ until only one mass shift remains in the list. Finally, the last remaining mass shift is added the result list $R$.

### Figures

**Algorithm 1** **Greedy algorithm for removing duplicated mass shifts**

**Input** $L$: A list of mass shifts sorted in the increasing order of the left boundary.

**Output** $R$: A list of non-overlapping mass shifts.

**while** $L$ contains $\geq 2$ mass shifts **do**
    Let $S_1$, $S_2$ be the first two mass shifts in $L$, and $(a_1,b_1)$,$(a_2,b_2)$ be
    the boundaries of $S_1$ and $S_2$, respectively
    **if** $b_1 \leq a_2$ **then** # Case 1: $S_1$ and $S_2$ do not overlap
        **remove** $S_1$ from $L$ and **add** $S_1$ to $R$
    **else if** $a_2 < b_1 < b_2$ **then** # Case 2: $S_1$ and $S_2$ partially overlap
        **remove** $S_2$ from $L$
    **else** # Case 3: $S_1$ fully covers $S_2$
        **remove** $S_1$ from $L$
    **end if**
**end while**
**add** the last mass shift in $L$ to $R$
**return** $R$

**Figure S1.** A greedy algorithm for removing duplicated mass shifts

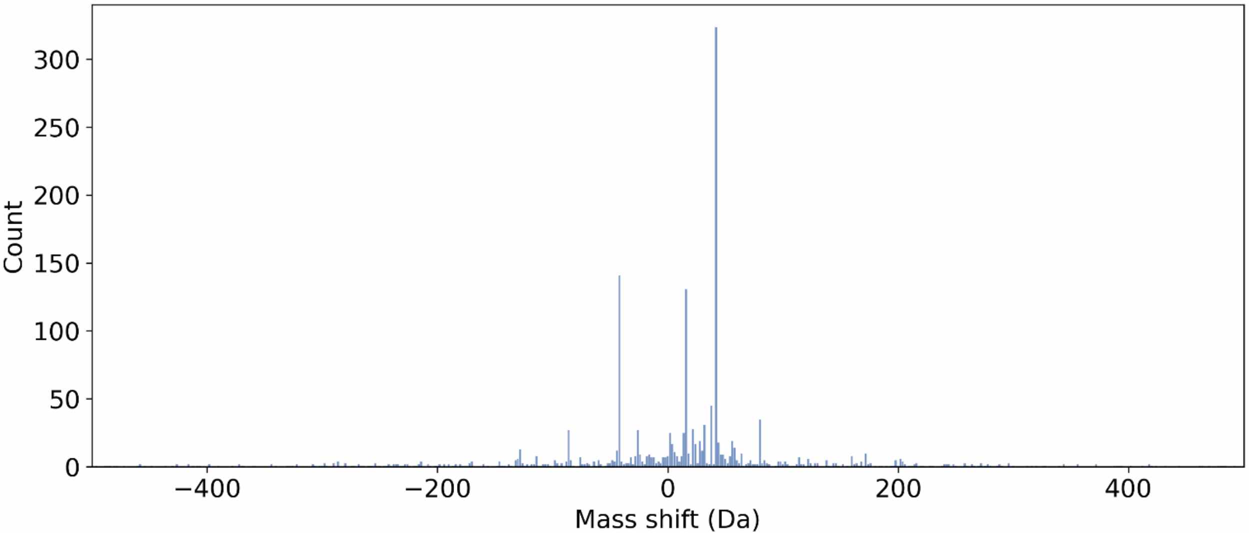


**Figure S2.** A histogram of mass shifts reported by TopPIC from the top-down MS data in the range [-500, 500] Da.

**Tables**

**Table S1.** High frequency PTMs reported by MSFragger in database search. Commonly modified residues and uncommonly modified residues are obtained from the UNIMOD database.

| PSI-MS Name | Description | Modified residues (common) | Modified residues (uncommon) | Mass shift (Da) | % PSMs |
|---|---|---|---|---|---|
| Carbamyl | Carbamylation | K | RCMSTY | 43.005814 | 1.10 |
| Cation: Na | Sodium adduct | DE | - | 21.981943 | 0.96 |
| Cation: K | Replacement of proton by potassium | - | ED | 37.955882 | 0.68 |
| Phospho | Phosphorylation | TSY | DHCRKE | 79.966331 | 0.58 |
| AEBS | Aminoethylbenzene-sulfonylation | - | HKSY | 183.035399 | 0.37 |
| Cation: Fe [III] | Replacement of 3 protons by iron | - | DE | 52.911464 | 0.34 |
| Methyl | Methylation | - | CHKNQRILEDST | 14.01565 | 0.28 |
| Cation: Fe [II] | Replacement of 2 protons by iron | - | DE | 53.919289 | 0.22 |
| Dimethyl | di-Methylation | - | KRNP | 28.0313 | 0.22 |
| Acetyl | Acetylation | K | CSTYHR | 42.010565 | 0.19 |
| Oxidation | Oxidation or Hydroxylation | MWH | DKNPFYRCGUEILQSTV | 15.994915 | 0.16 |
